# Polygenetic Determinants of Azole Resistance, Tolerance, and Heteroresistance in *Candida albicans*

**DOI:** 10.1101/2025.06.29.662174

**Authors:** Kyle S. Schutz, Cecile Gautier, Corinne Maufrais, Natacha Sertour, Marie-Elisabeth Bougnoux, Christophe d’Enfert, Luke M. Evans, Stacey D. Smith, Iuliana V. Ene

## Abstract

Azole antifungals are widely used to treat *Candida* infections, yet therapeutic failures are common. In addition to resistance, fungal populations can exhibit tolerance and heteroresistance, subpopulation-driven drug responses that can contribute to treatment failure. Here, we performed a genome-wide association study (GWAS) on 557 genetically diverse *C. albicans* isolates to identify genetic loci associated with fluconazole susceptibility, tolerance, and heteroresistance. We uncovered a complex, polygenic architecture underlying all three traits, involving novel loci linked to stress responses, cell cycle, and genome integrity. Notably, canonical resistance genes were absent from our top associations, reflecting either their low frequency or lineage specificity. Functional validation of several candidate genes confirmed distinct genetic determinants for each drug response. Heritability estimates and conditional analyses revealed that susceptibility and heteroresistance are governed by partially overlapping but largely independent loci, with some genetic variants displaying opposing effects on different traits. This limited overlap suggests that each phenotype represents a distinct evolutionary strategy for drug evasion, which may not be simultaneously addressed by a single therapeutic approach. Our findings underscore the polygenic nature of adaptation to antifungal drugs and expand our understanding of drug response mechanisms in *C. albicans*.

**SIGNIFICANCE:** Despite advances in antifungal therapies, treatment failures in *Candida albicans* infections remain common and poorly understood. Standard clinical diagnostic tools rely on susceptibility testing to predict treatment outcomes, yet many treatment-refractory infections are caused by strains that appear susceptible in laboratory tests. This study provides the first genome-wide dissection of three distinct drug responses – susceptibility, tolerance, and heteroresistance - across a global population of *C. albicans* isolates. By identifying novel genetic contributors beyond canonical resistance genes, our findings shift the current paradigm toward a more nuanced, polygenic model of antifungal adaptation. These results underscore the importance of integrating tolerance and heteroresistance into clinical and research frameworks and point to new molecular pathways that could be leveraged to enhance antifungal strategies.

## INTRODUCTION

Treating human fungal infections remains a major challenge because fungi share many cellular processes with humans, leaving few drugs that are both effective and safe (1). In addition to limited treatment options, emerging drug resistance further narrows treatment options. Resistance, or the ability to grow despite drug treatment, is routinely tested through susceptibility assays to guide clinical decisions. Among fungal pathogens, the diploid yeast *Candida albicans* is one of the most frequently isolated species, and a pathogen of critical priority, as classified by the World Health Organization (2). Resistance to azoles, a first-line treatment against diverse fungi, is rapidly increasing in *Candida* species (3). Azoles are fungistatic drugs that inhibit growth by binding to lanosterol 14-alpha-demethylase (encoded by the *ERG11* gene), thereby disrupting ergosterol synthesis, a major component of fungal membrane, weakening the yeast cell, and enabling the immune system to clear the infection (4). Azole-resistant phenotypes can arise in *C. albicans* through multiple mechanisms, including via mutations in the gene encoding the drug target (*ERG11*) or through the upregulation of efflux pump genes (e.g., *CDR1*, *CDR2*, *MDR1*), which reduces intracellular drug accumulation (4, 5). Although invaluable as a clinical assay, antifungal susceptibility testing alone does not capture the full spectrum of fungal azole responses. The rising incidence of azole treatment failure in *C. albicans* infections, even among susceptible isolates, indicates that this pathogen employs diverse and often poorly understood mechanisms to evade the effects of antifungal treatment (6).

Azole tolerance and heteroresistance have been previously described in *C. albicans* and are hypothesized to enable persistent infections. Tolerance refers to the ability of fungal cells to grow slowly above the minimum inhibitory concentration (MIC). Unlike susceptibility, tolerance is independent of drug concentrations and relies on the functioning of stress response pathways such as the calcium-calcineurin pathway and the heat shock response (7). Presumably, cell-to-cell variation in the activity of stress pathways enables different tolerance levels (7). Consistent with this, inhibition of stress response pathways can reduce tolerance to fluconazole, thereby enhancing drug efficacy without altering susceptibility (6, 7). Tolerant strains were more likely to persist in human infections than non-tolerant isolates (7).

Distinct from tolerance and resistance, heteroresistance involves varying antimicrobial susceptibility within a clonal population, and can serve as a bet-hedging strategy to increase population fitness in the presence of a drug. Specifically, rare, highly resistant subpopulations exist within an otherwise susceptible population. Such subpopulations can expand under drug pressure, leading to population rebound (8, 9, 10). Standard susceptibility assays cannot detect heteroresistant populations, which require quantitative assessment via drug-gradient plating and growth fraction analysis (11). Heteroresistance has been recently described in fungi, including *Cryptococcus neoformans* and several *Candida* species (10, 11, 12). Here, it was associated with copy number variations of beneficial genes (10, 13) or increased drug efflux via upregulation of ABC drug transport genes (12). Heteroresistant isolates can persist in azole-treated mice, suggesting that heteroresistance, like tolerance, contributes to treatment failure and the emergence of full resistance (12).

In *C. albicans*, the genetic mechanisms underlying susceptibility, tolerance, and heteroresistance, may reflect evolutionary dynamics that span microbe-microbe interactions in natural environments as well as a long-term association with humans. Resistance often arises via novel advantageous large-effect mutations that sweep to fixation across the entire population, as seen in diverse pathogens both within (14, 15, 16) and outside the fungal kingdom (17). Moreover, given the constraints imposed by pleiotropic effects, these mutations may be concentrated in a few ‘hotspot’ loci, which are repeatedly co-opted during independent gains of resistance (16, 18). By contrast, tolerance and heteroresistance are phenomena wherein only a subpopulation overcomes the drug exposure. In this case, multiple advantageous variants may persist and compete before one ultimately dominates (14, 19). While less studied than drug resistance, tolerance and heteroresistance likely depend on small-effect and trade-off mutations that rarely become fixed in populations (20, 21, 22).

Genome-wide association studies (GWAS) have been used to dissect genotype-phenotype relationships for two decades and are a powerful tool for characterizing the genetic architecture of complex phenotypes (23). GWAS applications in fungal biology remain limited, primarily exploring pathogenicity and antifungal resistance mechanisms. Examples of fungal GWAS studies include studying communication in *Neurospora crassa* (24) and itraconazole sensitivity in *Aspergillus fumigatus* (25). In *Candida*, one GWAS study examined azole resistance across six species (26), while another analyzed *C. auris* resistance to nine antifungals (27). Both uncovered associations with expected genes (e.g., *ERG11*), consistent with the repeated recruitment of major effect loci, as well as a range of novel resistance-associated loci (26, 27). Despite the clinical importance of *C. albicans*, GWAS approaches have not yet been applied to study its drug response genetics. Furthermore, population-level fungal genomic studies have overlooked tolerance and heteroresistance mechanisms.

Here, we employ GWAS to investigate the genetic basis of azole response variation in the model fungal pathogen *C. albicans* by leveraging a collection of 557 human and environmental isolates. GWAS revealed variants associated with fluconazole resistance, but also with tolerance, and heteroresistance, which has not been previously done in any fungal species. We hypothesized that resistance would involve large-effect loci (14, 16, 18) and may be linked to genomic ‘hotspots’ (28, 29). Given that, in other fungi, heteroresistance involves some of the same mechanisms that enable resistance (e.g., increased dosage of drug efflux pumps, increased copies of drug target), we expected an overlapping genetic basis. By contrast, tolerance operates through independent stress-related pathways and varies quantitatively across strains (7), suggesting a distinct and more complex genetic architecture with many contributing single nucleotide polymorphisms (SNPs) explaining a smaller proportion of the variance. Using a mixed-linear model that accounts for population structure and relatedness among isolates, we identified 21 loci under stringent thresholds and 162 loci under relaxed thresholds that extend beyond the canonical resistance mechanisms described for *C. albicans*. We identified novel loci linked to susceptibility involved in DNA repair and cell wall integrity. In contrast, heteroresistance was primarily associated with loci involved in DNA replication and genome stability. Interestingly, tolerance was associated with loci related to lipid homeostasis, metabolism, stress responses, and DNA replication, further emphasizing the complexity of these adaptive mechanisms. Our GWAS analysis also suggests that genetic factors influencing azole response may be interconnected, potentially leading to trade-offs among these traits. Many of the loci associated with the three phenotypes have not been previously implicated in adaptation to azoles, suggesting potential new drug targets that could serve the growing need for antifungal strategies.

## RESULTS

### *C. albicans* isolates display diverse fluconazole responses that span genetic clusters

Phenotypic analysis of 557 diverse clinical and environmental *C. albicans* isolates revealed pervasive yet variable susceptibility, tolerance, and heteroresistance across different genetic lineages (Fig. 1A and B, Fig. S1A). For example, 59.78% (333/557) of the isolates were susceptible to fluconazole (S, MIC<u>_50_</u> <4 μg/ml), and among those resistant, 12.75% (71/557) had MIC_50_ levels ≥256 μg/ml FLC. For isolates where tolerance and heteroresistance was detected, high tolerance (SMG ≥0.5) was observed in 75.67% (367/486) of isolates, and 55.45% (234/424) exhibited high heteroresistance, with ≥5% of the population displaying elevated resistance relative to the remainder (Fig. 1A and B; Fig. S1B, Table S2). Resistance, high tolerance, and high heteroresistance were found not only in the 530 human-associated isolates but also in strains isolated from environmental sources. Among the 16 isolates from food spoilage and three from starlings, 2 were resistant, 14 were highly tolerant, and 12 were highly heteroresistant (Table S2). Thus, strains capable of withstanding azole treatment can also be isolated outside of the human host.

**Figure 1.**
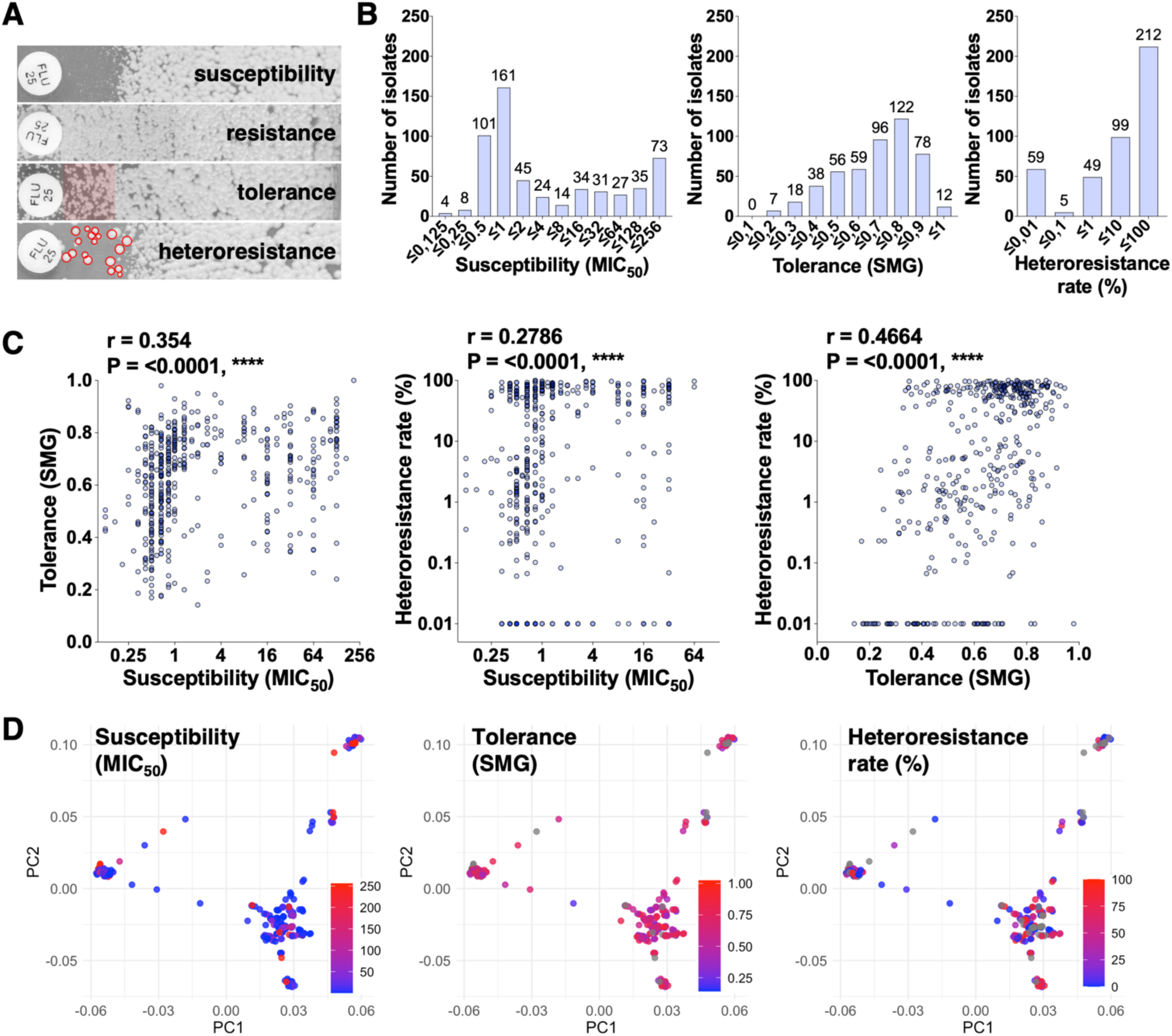
Phenotypic diversity of azole responses across the 557 *C. albicans* isolates. (A) Representative disk diffusion assays (fluconazole, 25 μg) showing growth patterns for susceptible (no growth), resistant (confluent growth), tolerant (delayed growth at 48 h, shaded area), and heteroresistant (isolated colonies within inhibition zone, red circles) isolates. (B) Distribution of susceptibility (MIC50), tolerance (supra-MIC growth, SMG), and heteroresistance rates (%) across 557 isolates. Susceptibility is scored as the minimum inhibitory concentration that reduces growth by 50% (MIC_50_); tolerance is measured as the growth at drug concentrations higher than the MIC_50_ (SMG) relative to the growth in the absence of drug at 48 h; heteroresistance is quantified as the number of non-confluent colonies at 128 μg/mL FLC divided by the number in the absence of FLC (short-PAP assay (11)). (C) Pairwise phenotypic correlations between each two drug responses (Spearman r, **** *P* < 0.0001). Each point represents a single isolate. (D) PCA of SNP data estimated from the genome-wide relationship matrix (GRM) with isolates colored by phenotype values for susceptibility, tolerance, and heteroresistance, overlaid on ML phylogeny clusters. Each point represents a single isolate.

These three drug responses appeared broadly positively correlated, although large numbers of isolates showed contrasting phenotypes. For instance, susceptibility and tolerance were significantly correlated (r = 0.354, *P* <0.001, nonparametric Spearman correlation, Fig. 1C), yet among 152 highly resistant isolates where tolerance was detected, 129 (84.87%) exhibited high tolerance, while 23 (15.13%) showed low tolerance (Fig. S1C). Similarly, among the 333 susceptible isolates, 238 (71.47%) were highly tolerant, while 95 (28.53%) had low tolerance (Fig. S1C). Similar moderate but significant correlations were found between susceptibility/tolerance and heteroresistance (*P* <0.001, nonparametric Spearman correlations, Fig. 1C), although the set of isolates without heteroresistance showed a wide range of susceptibility and tolerance values. Importantly, many susceptible strains (86.2%, 287/333) expressed some degree of heteroresistance, either at low (35.1%, 117/333) or high (51.1%, 170/333) rates (Fig. S1C). The presence of strains with diverse drug responses suggests that these phenotypes are controlled, at least in part, by genetically distinct mechanisms.

Next, we examined the distribution of azole responses across lineages to determine whether each is broadly distributed across the isolates or confined to specific clades, which could potentially confound association results. A principal component analysis (PCA) of the 145,129 SNPs in our dataset recovered several distinct clusters, consistent with strong genetic structure within *C. albicans* (30). Highly resistant phenotypes were common within each cluster, and similar distributions were observed for tolerance and heteroresistance (Fig. 1D). We also examined the distribution of the drug response phenotypes across the *C. albicans* clades (30) by estimating a maximum likelihood (ML) phylogeny from the SNP dataset. Each clade presented a range of drug response phenotypes and accordingly, was widely convergent across the tree. For example, highly heteroresistant strains were present in all 17 clades (Fig. S2). The wide phylogenetic distribution of isolates sharing the same phenotype provides a strong foundation for using GWAS to identify shared genetic architecture.

### The isolate collection exhibits extensive variation in chromosome copy numbers and SNP frequencies

Whole genome sequencing of the 557 isolates revealed diversity in both SNPs and copy number variations (CNVs), highlighting key patterns in chromosomal architecture and variant distribution, which provide context for interpreting the GWAS findings. The SNP distribution patterns revealed genetic diversity with important functional implications. Chr6 exhibited the highest SNP density across isolates (average of 11.1 SNPs/kbp versus 8.5 SNPs/kbp across the genome), followed by Chr5 (9.9 SNPs/kbp), while Chr3 showed the lowest polymorphism levels (6.9 SNPs/kbp, Fig. S3A and B). This non-random genomic distribution suggests chromosome-specific evolutionary constraints, where elevated SNP densities in Chr5 and Chr6 may reflect adaptive diversification. Changes in chromosome copy numbers, most frequently trisomies (3 instead of 2 copies), were observed across multiple chromosomes, with Chr5 and Chr7 showing the highest variability (Fig. S3C and D). This aligns with observations of frequent aneuploidy among drug-resistant, drug-tolerant, and host-adapted *C. albicans* isolates (31, 32). Only one isolate displayed monosomy (of Chr 4, 6, and 7), suggesting that selection favors increased rather than decreased gene dosage.

We also noted significant phenotypic differences between aneuploid and euploid isolates. Aneuploid strains displayed higher resistance, with 53.8% of aneuploids showing MIC_50_ ≥32 μg/mL compared to only 20.3% of euploids (Fig. S4A). Similarly, tolerance was higher in aneuploids (60% of aneuploids with SMG ≥0.7 vs 41.7% of euploids), as was heteroresistance (64.5% vs 48.9% with rates ≥10%). Notably, while aneuploidy was associated with increased resistance (mean MIC_50_ of 88.8 for aneuploids vs 41.5 for euploids) and higher heteroresistance rates (39.4% vs 32.6%), we observed no specific association between these phenotypes and particular aneuploid chromosomes (Fig. S4B and C). These findings suggest that aneuploidy itself, rather than gains of specific chromosomes, may enhance fitness under azole stress.

### Associated loci suggest alternate genetic mechanisms involved in azole drug adaptation

To characterize the genetic basis of each phenotype, we first examined its genetic architecture through GWAS. We recovered a total of 21 loci under stringent thresholds and 162 under relaxed thresholds associated with the three drug responses (see Materials and Methods for stringent and relaxed α thresholds; Fig. 2, Table S3, Fig. S5). Among stringent loci associated with susceptibility (n=9), several were located within functionally annotated genes, including *EXO1*, *MRV1*, and *orf9.952* (Fig. 2A and 3A). Based on orthology with *S. cerevisiae*, *EXO1* could contribute to DNA mismatch repair and homologous recombination (33), while the other two genes lack predicted functions. We also identified three significantly associated variants in *RLM1*, a transcription factor involved in cell wall stress response (34). Interestingly, 7/9 stringent and 31/44 relaxed loci among the susceptibility variants were located on Chr5. Additional resistance-associated loci were identified in genes implicated in lipid homeostasis (*DNF2*), membrane trafficking (*RCY1*), cell wall integrity (*INP54*, *KRE62*), and DNA replication and repair (*RAD57*, *RFC3*) (Table S3). This is in line with the importance of the plasma membrane and cell wall in maintaining cell homeostasis during azole treatment (4, 35).

**Figure 2.**
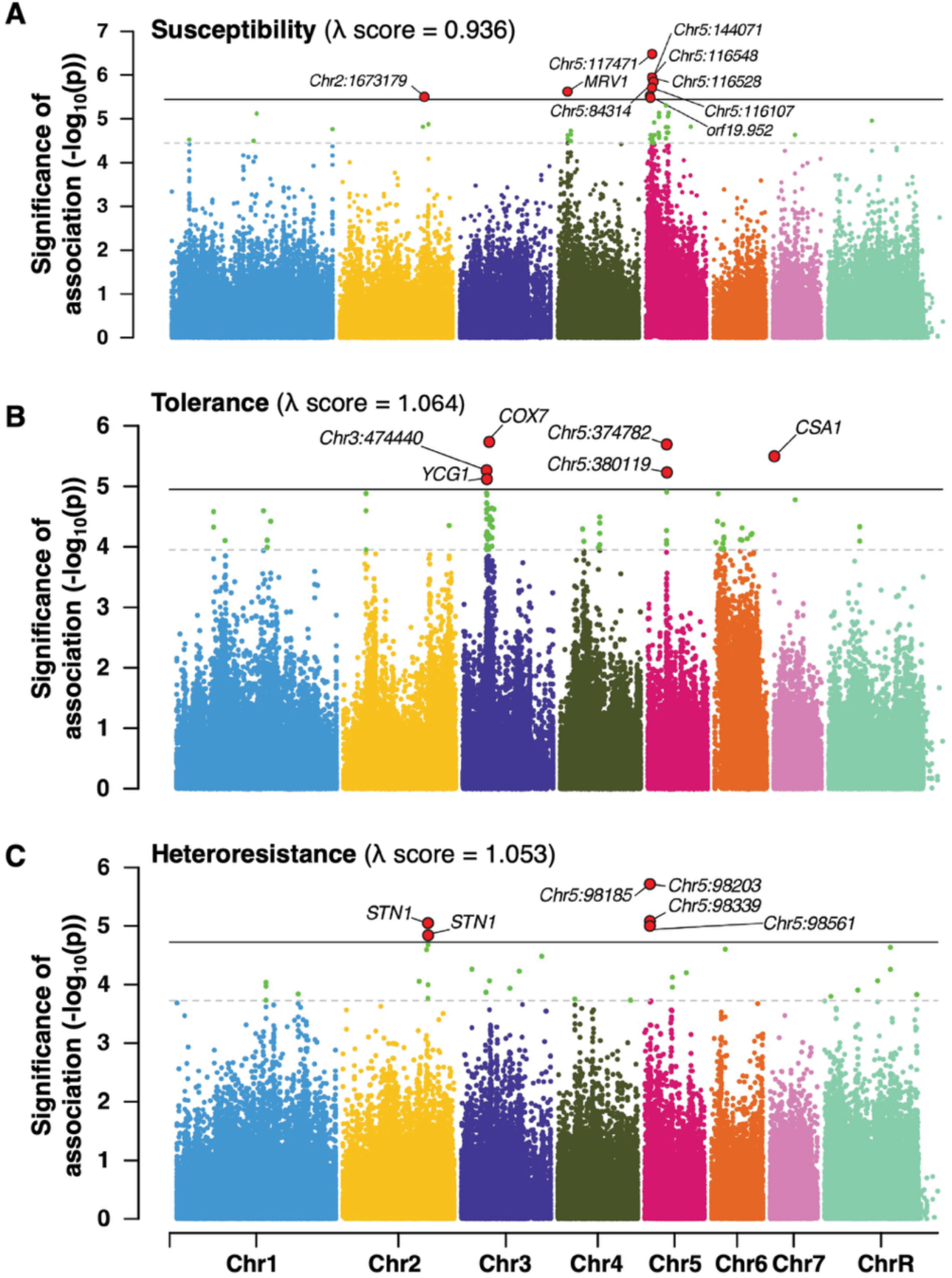
Distinct genetic architectures of azole response traits. Manhattan plots for (A) susceptibility, (B) tolerance, and (C) heteroresistance. The y-axis shows the –log₁₀ of the *P* values from each GWAS, representing the statistical significance of association between each SNP and the phenotype of interest. Higher -log₁₀(p) values indicate a stronger association. Dashed lines indicate significance thresholds (top: stringent; bottom: relaxed, as determined by power analyses for each phenotype (see Materials and Methods). *C. albicans* chromosomes are labeled (Chr1–7, ChrR) on the x-axis. A λ score was calculated for each GWAS to assess the performance of the MLMA model given the data. A λ score of 1 indicates that the test statistics are not inflated due to cryptic relatedness or population structure in the sample.

**Figure 3.**
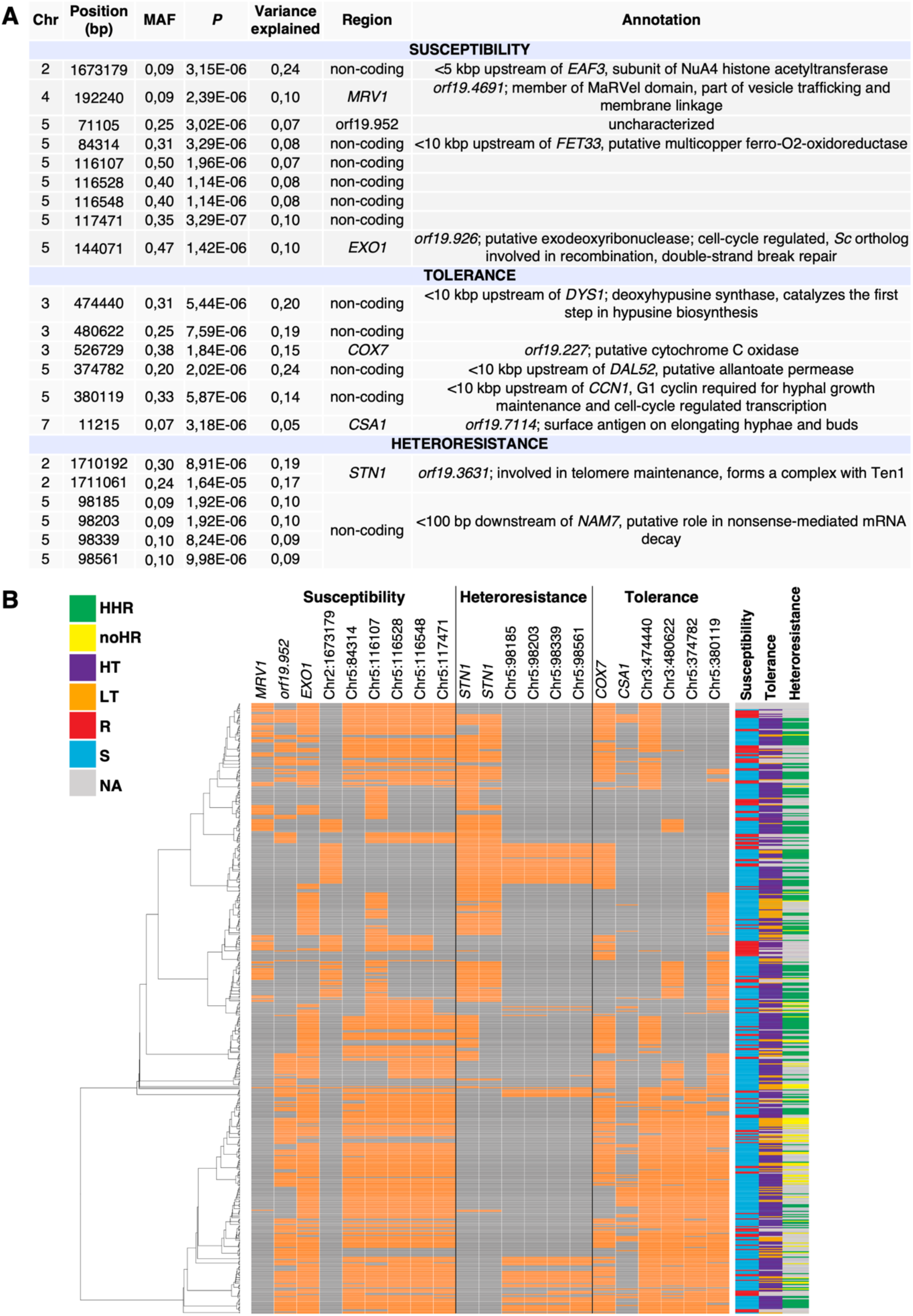
Candidate loci associated with azole response phenotypes. GWAS summary statistics for stringent associations for (A) susceptibility, (B) tolerance, and (C) heteroresistance, annotated by genomic position, minor allele frequency (MAF), *P* value, proportion of variance explained by SNP (PVE), genomic region affected, and gene function according to the *Candida* Genome Database (Skrzypek et al., 2017). (B) Phylogenetic distribution of independently associated significant loci by phenotype. Carriers of significant loci (either heterozygous or homozygous) identified in the GWAS were mapped onto the estimated maximum likelihood (ML) phylogeny, with variant presence/absence displayed as a heatmap. Susceptibility, tolerance, and heteroresistance levels for each isolate are also included.

Like resistance, several heteroresistance-associated variants were located on Chr5 (4/6 stringent loci) (Fig. 2C and 3A). Interestingly, we recovered variants in genes with known or predicted functions in telomere protection and chromosome stability (*STN1)* (36), actin organization (*ARC40*), nuclear import (*MTR10*), endoplasmic reticulum unfolded protein response (*SKI2*), and DNA replication (*orf19.2796*, an ortholog of the polymerase gene Sc*POL12*), indicating the importance of genome stability in heteroresistance (Fig. 3A, Table S3).

By contrast, tolerance phenotypes were less influenced by variants on Chr5, with only 7.3% (7/96) of significantly associated variants across both thresholds found here (Table S3). Instead, 42.7% (41/96) and 18.8% (18/96) of these variants were located on Chr3 and Chr6, respectively (Fig. 2B, 3A, Table S3). Several tolerance-associated variants were located in genes with known or predicted roles in lipid homeostasis (*PLB5*, *SEC59*), metabolism (*ADO1*, *SAP1*, *DUR1,2*, *LAP41*, *orf19.1626*), ion homeostasis (*ALR1)*, and stress responses (*CAP1, orf19.3051*) (37-40). We also recovered loci associated with DNA replication/chromosome segregation (*YCG1, BFR1, LRP1, orf19.5701*), highlighting the potential role for aneuploidy in tolerance. The most significantly associated variant for tolerance was a missense mutation in *COX7*, encoding a cytochrome C oxidase; its *S. cerevisiae* ortholog influences mitochondrial function and triggers compensatory stress pathways (41). Our GWAS results for tolerance in *C. albicans* indicate multiple genetic pathways and stress response mechanisms, contrasting with the genetic architectures seen in resistance and heteroresistance.

Overall, GWAS revealed that fluconazole responses in *C. albicans* are associated with distinct but overlapping genetic architectures, with susceptibility and heteroresistance enriched for variants on Chr5 and genes involved in DNA repair and stress responses, while tolerance appears influenced by diverse loci, particularly on Chr3 and Chr6, implicating pathways related to lipid homeostasis, metabolism, stress responses, aneuploidy, and mitochondrial function.

### Narrow phylogenetic distribution or rarity of canonical resistance mechanisms limits their detection in GWAS

Notably, we did not recover significantly associated variants in any of the canonical genes that mediate azole resistance in *C. albicans*, including the drug target *ERG11*, the ergosterol regulator *UPC2*, the efflux pumps *CDR1*, *CDR2*, *MDR1*, or their transcriptional regulators *TAC1* and *MRR1* (Lee et al. 2023). Other GWAS studies on antifungal resistance have similarly failed to identify significantly associated loci within these known genetic drivers (26). On one hand, the most informative SNPs in such analyses are typically those weakly structured by phylogeny but strongly linked to the phenotype. For example, canonical genes like *MRR1* and *MDR1* have a high number of variants (57 and 29, respectively) that are broadly distributed across the phylogeny, yet these variants also failed to emerge as significantly associated through GWAS (Fig. S6). Conversely, rare variants were excluded during quality control filtering for minor allele frequency (MAF), retaining only those present in at least 5% of the population (see Materials and Methods). In our sampled population, we identified 81 SNPs in *ERG11* from the unfiltered SNP dataset. Of these, 76 were excluded from the GWAS due to low MAF, while 5 were fixed within specific subpopulations (Fig. S6). None of the 5 variants confer resistance (42, 43). These examples suggest that canonical resistance mechanisms, while well-characterized in clinical isolates, may not be consistently predictive of resistance phenotypes across genetically diverse backgrounds.

### Conditional and joint analysis identifies independent loci driving drug response variation

Through GWAS, we identified two regions displaying a high density of phenotype-associated loci: Chr3 (450-500 kbp) that included 24/96 tolerance loci (both stringent and relaxed), and Chr5 (93–160 kbp) that included 14 resistance and 4 heteroresistance loci (Table S3). Given the concentration of signals in these regions, we performed a conditional and joint analysis (COJO) using GCTA to determine whether any of the significantly associated variants represent independent associations or a single signal for each phenotype (44). This type of analysis can also identify secondary association signals at a locus by disentangling independent signals in regions of high linkage disequilibrium (LD) (44). COJO identified three independent tolerance associations on Chr3: 473398, 474440, and 480622 (Fig. 4A, Table S4). Chr3:473398 maps to *DYS1*, which encodes deoxyhypusine synthase. Its *S. cerevisiae* ortholog is essential, and its depletion results in G1 cell cycle arrest (45). Both Chr3:474440 and Chr3:480622 are intergenic, but the latter lies between the *YCG1* and *CAP1* genes. Independent variants in these loci may implicate both Cap1, a transcription factor regulating the oxidative stress response, and Ycg1, a putative condensin involved in chromosome segregation in *S. cerevisiae* (46, 47). However, fine mapping and GWAS integration with eQTL data will be needed to inform the role of these genes in the phenotype. For the Chr5 region, COJO identified Chr5:117471 and Chr5:155398 as independent associations for susceptibility, and Chr5:98185 and Chr5:122508 for heteroresistance (Fig. 4B, Table S4). Chr5:117471 and Chr5:98185 represent GWAS-identified intergenic SNPs for susceptibility and heteroresistance, respectively (Table S3). The newly identified association at Chr5:122508 lies within *UBC13*, which encodes a predicted ubiquitin-conjugating enzyme potentially involved in the DNA damage response. In addition, Chr5:155398 maps to an intergenic region between *HMS1* and *RMT2,* encoding a transcription factor of morphogenesis, and an arginine methyltransferase, respectively (48, 49).

**Figure 4.**
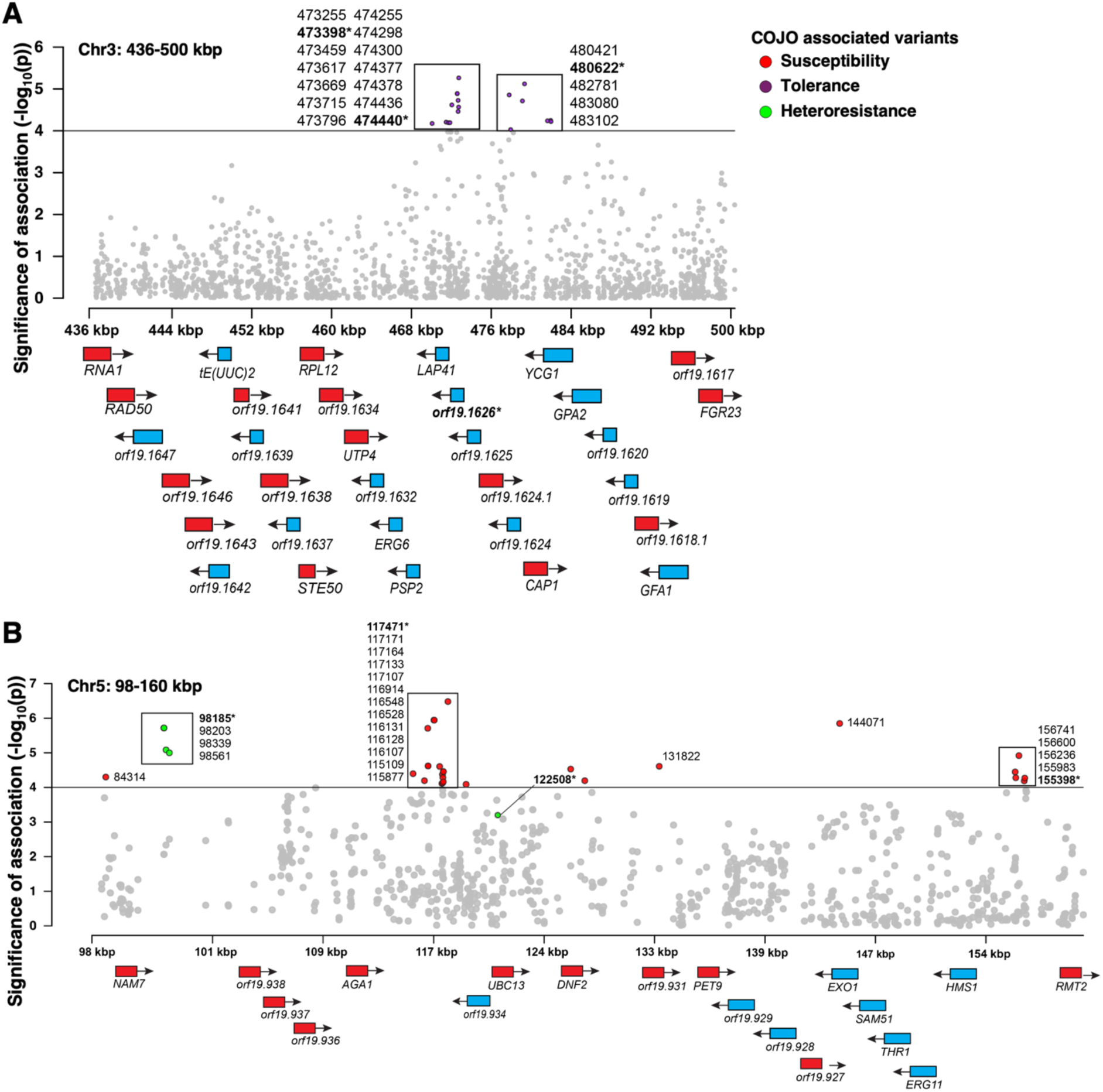
Conditional and joint (COJO) analyses of genetic regions harboring multiple significant loci. (A) An overlapped Manhattan Plot of SNP associations from GWAS for tolerance on Chr3. Purple dots represent significant loci for tolerance. Boxes indicate SNPs in high LD (r^2^ ≥0.80) with the bolded and asterisked SNPs being the true associations recovered from COJO. (B) An overlapped Manhattan Plot of SNP associations from GWAS for susceptibility and heteroresistance on Chr5. Red and green dots represent significant loci for susceptibility and heteroresistance, respectively. Boxes indicate SNPs in high LD (r^2^ ≥0.80) with the bolded and asterisked SNPs being the true associations recovered from COJO.

These findings reveal that multiple independent genetic variants underlie the clustered GWAS signals on Chr3 and Chr5, highlighting the complex and polygenic nature of azole responses. COJO analysis in these regions identified several conditionally independent associations, reflecting distinct genetic effects. Although the COJO-retained variants are statistically independent, broader genomic regions surrounding these loci exhibit complex LD patterns (r² > 0.8), possibly due to rarer variants excluded during quality control (Fig. S7). Notably, five out of 7 associations were intragenic, suggesting that regulatory elements may contribute to the observed phenotypic variation.

### GWAS hits explain modest variance, but azole responses are highly heritable

The loci associated with susceptibility, tolerance, and heteroresistance (under both stringent and relaxed thresholds) together explained 6.36%, 14.59%, and 3.99% of the phenotypic variance, respectively (Table S3). These values represent the cumulative variance explained by individual GWAS-significant SNPs, providing a first approximation of how much the identified hits contribute to each trait. As phenotypic variance arises from genetic, environmental, and stochastic factors, GWAS-significant SNPs typically explain only a modest proportion of that variance. The 4–15% captured here aligns with expectations for complex, polygenic traits (50).

To more accurately capture the genetic architecture, including the effects of LD and potential polygenicity, we estimated SNP-based heritability (*h²*_SNP_). Since such estimates may include residual LD between SNPs and inflate the total variance explained, we partitioned the phenotypic variance into two components: (1) variance explained by genome-wide SNPs using a GREML (Genomic Restricted Maximum Likelihood) analysis, and (2) residual variance (51, 52). We found that tolerance had the highest ratio of genetic variance to phenotypic variance (*h²*_SNP_, 76.3%), followed by susceptibility (72.1%) and heteroresistance (66.5%) (Table S5). This pattern suggests that each phenotype is influenced by numerous additive common variants. The high heritability estimates across all three traits point to a polygenic architecture, with substantial contributions from common SNPs. Notably, these heritability estimates exceed the variance explained by the GWAS significant loci identified here, underscoring that larger sample sizes will likely reveal additional associated loci. For each phenotype, we observed moderate residual variance (23.7% to 33.5%), suggesting that non-genetic factors are still contributing (Table S5).

We then partitioned the estimated *h²*_SNP_ from the GRMEL by chromosome to examine how genetic contributions are distributed across the genome (Figure S8A-C). This approach allowed us to test whether the variance explained scales with chromosome size, as predicted under a highly polygenic model where causal variants are broadly distributed (53). Across all three phenotypes, we observed a weak, non-significant, positive correlation between chromosome length and SNP-heritability. This suggests that while larger chromosomes may contribute more to the genetic variance, the distribution of genetic effects is likely more complex and not solely dependent on chromosome size.

### Antagonistic genetic effects between susceptibility and heteroresistance suggest evolutionary trade-offs

To further investigate the shared genetic basis among these traits, we used a bivariate GREML model to estimate the genetic correlation (*rG*) between each pair of phenotypes (Table S5 and Fig. S8D) (54). The strongest correlation was observed between susceptibility and tolerance (*rG* = 0.428, *P* = 0.006), consistent with phenotypic trends showing that tolerant isolates are often resistant (Fig. S1C). In contrast, tolerance and heteroresistance had a weak and non-significant genetic correlation (*rG* = 0.175, *P* = 0.3620), suggesting distinct genetic drivers. Interestingly, we observed a negative genetic correlation between susceptibility and heteroresistance (*rG* = –0.404, *P* = 0.0105), indicating that alleles conferring resistance may antagonize heteroresistance. This antagonism is further illustrated by a high SNP-density region on Chr5 identified in our GWAS (Fig. 2), where SNPs in high LD were associated with increased heteroresistance but decreased resistance (Table S6). For example, a SNP in the gene *STN1*, which encodes a protein involved in telomere maintenance, exhibited opposing effects: the effect size was positive for heteroresistance (24.07), but negative for susceptibility (–7.84). This suggests that increased genomic instability might facilitate the formation of heteroresistant subpopulations (e.g., through aneuploidy) while antagonizing pre-existing resistant genomic configurations. Thus, while some traits share overlapping genetic influences, others exhibit antagonistic relationships, underscoring the complexity of azole response phenotypes and the potential trade-offs in targeting specific resistance mechanisms.

### Loci identified through GWAS map to genes implicated in azole responses

Given the high number of variants associated with coding regions identified in our GWAS and COJO analyses, we aimed to determine whether altering these genes affects azole responses. We characterized knockout (KO) and overexpression (OE) mutants in the SC5314 reference strain, using full gene deletions or tetracycline-inducible expression (55). The susceptibility assays revealed gene-specific effects, with several mutants displaying altered MIC_50_ values compared to the corresponding wild type (parent) strain (Fig. 5A). Notably, the deletion of *RCY1* increased susceptibility >2-fold, while the deletion and the overexpression of *RLM1* altered MIC_50_ levels in opposite directions, establishing both genes as regulators of susceptibility. Rlm1 is a transcription factor involved in cell wall integrity (34), while Rcy1 is an F-box protein involved in endocytic membrane traffic (56). The alteration of several tolerance-associated genes affected susceptibility, including *ADO1*, *CAP1*, *LRP1*, *orf19.1611*, *orf19.3051*, and *SAP1* (Fig. 5B). However, deletion of *orf19.1625*, an uncharacterized gene, reduced tolerance 0.44-fold while leaving susceptibility unchanged, demonstrating that these phenotypes can be genetically uncoupled (Fig. 5B). These results demonstrate the value of population genetics in uncovering cryptic susceptibility and tolerance determinants that would have been missed by candidate gene approaches.

**Figure 5.**
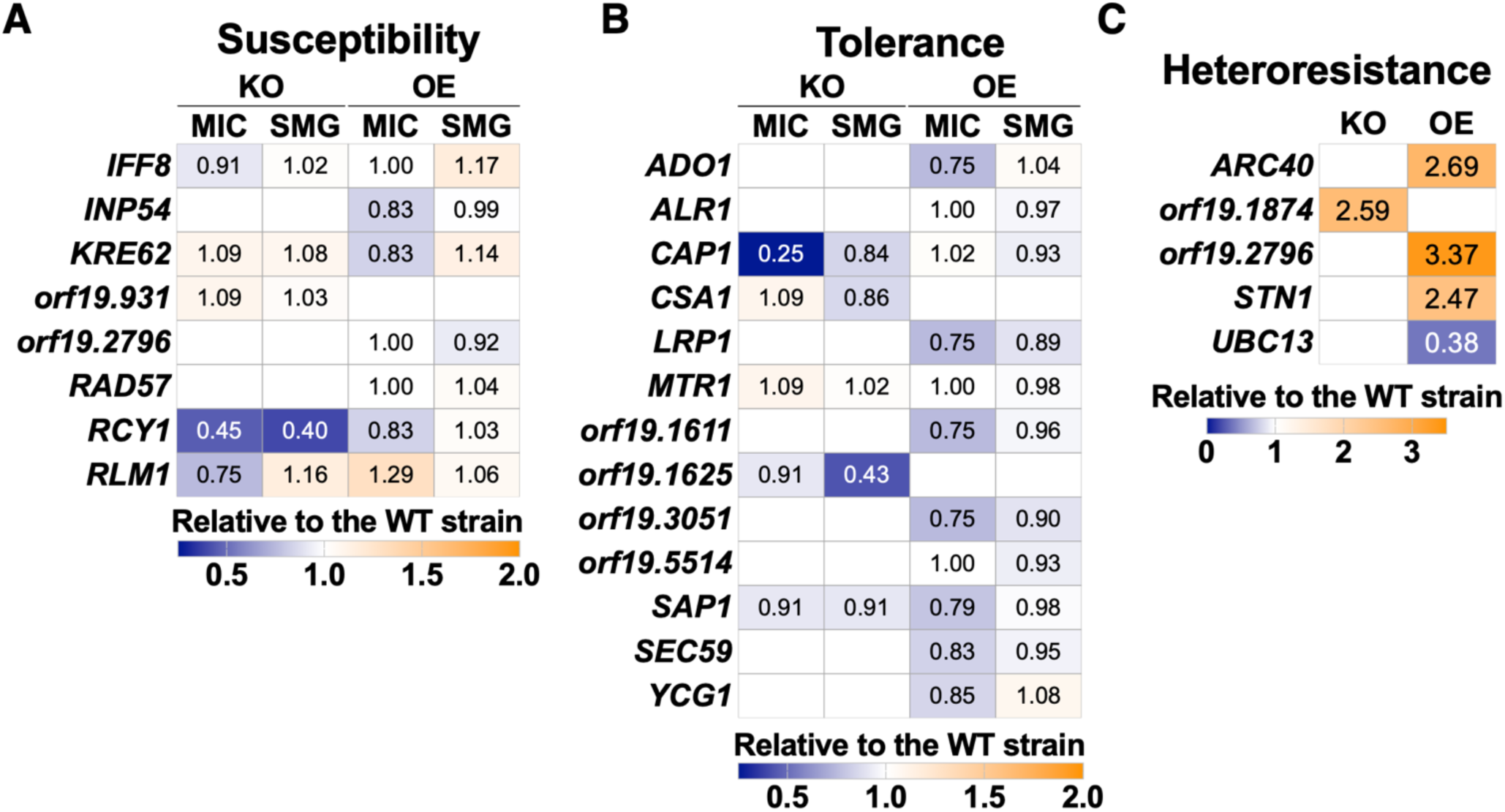
Functional validation of candidate genes harboring significantly associated GWAS loci. Relative phenotypic changes (vs. corresponding wild type (WT) isolates) in (A) susceptibility (MIC_50_), (B) tolerance (SMG), and (C) heteroresistance in knockout (KO) and overexpression (OE) strains.

The testing of heteroresistance-associated genes revealed the most pronounced effects, with the deletion of *orf19.1874* and the overexpression of *ARC40*, *orf19.2796*, and *STN1* increasing heteroresistance rates (Fig. 5C). Stn1 is involved in telomere maintenance, while Arc40 is similar to *S. cerevisiae* Arc40, a protein involved in actin filament organization (36, 57). orf19.1874 is similar to *S. cerevisiae* Mek1, which suppresses double-strand break repair to promote DNA recombination, while orf19.2796 is orthologous to *S. cerevisiae* Pol12, a subunit of DNA polymerase (58). Furthermore, the overexpression of *UBC13*, a ubiquitin conjugating enzyme involved in DNA damage response (59), identified through COJO, decreased heteroresistance, indicating that an overactivated DNA damage response might limit the formation of heteroresistant cells (Fig. 5C). Importantly, mutants that altered heteroresistance did not affect susceptibility or tolerance (Table S7), further supporting that these phenotypes are genetically distinct and can be uncoupled. This functional analysis revealed that multiple genes recovered from our study impact azole responses. However, they are likely involved in a polygenic response to the presence of azoles, as captured by our GWAS.

## DISCUSSION

Effective treatment of *C. albicans* infections requires strategies that inhibit fungal growth while minimizing harm to the human host. Although azoles remain a frontline therapy, the rising rates of resistance and recalcitrant infections are narrowing treatment options (1, 60). How *C. albicans* escapes azole treatment represents a complex and context-dependent evolutionary response to drug pressure, shaped by population size, diverse genetic mechanisms, heritability, fitness trade-offs, and stochasticity (1). Like other pathogenic traits, drug adaptation can be fueled by remarkable genomic plasticity, enabling numerous genetic routes to phenotypic variation across populations (62). Our phenotypic analysis of 557 genetically diverse *C. albicans* isolates revealed that susceptibility, tolerance, and heteroresistance are neither mutually exclusive nor restricted to specific clades. Instead, these drug responses are dispersed throughout the phylogeny, complicating efforts to predict treatment efficacy based on evolutionary lineage alone (Fig. S2). This has important implications for antifungal drug development, emphasizing the need for treatments that remain effective across genetically diverse *C. albicans* populations, as expanded sampling will likely uncover additional loci contributing to these complex phenotypes.

Our GWAS identified associations in novel loci for each phenotype, which were outside of well-characterized azole resistance genes. Susceptibility associated primarily with loci involved in DNA repair, cell wall integrity, and membrane trafficking, many of which were concentrated on Chr5. Tolerance, in contrast, was linked to diverse loci, many located on Chr3 and Chr6, implicating pathways such as lipid homeostasis, mitochondrial function, and stress responses, reflecting its quantitative and complex nature. Heteroresistance showed unique associations with genes related to genome stability and DNA replication, suggesting a role for karyotypic variation in subpopulation-driven drug evasion. Notably, associations with *BFR1*, *EXO1*, *RAD57*, *RFC3*, *STN1,* and *YCG1* suggest that azole responses in *C. albicans* may be more tightly linked to DNA replication and genome stability than previously recognized. These diverse associations point to a highly polygenic architecture underlying these phenotypes. While our focus on common variants does not rule out the role of rare, lineage-specific mutations, our findings suggest that targeting conserved stress responses or replicative mechanisms may be more effective at the population level than focusing on individual genes. Many of these variants may be under strong lineage-specific selection or exhibit high phylogenetic signal, thereby confounding standard GWAS approaches. Furthermore, population-level variation may arise from distinct combinations of rare variants, epistatic interactions, or non-canonical pathways that are not captured in single-locus association tests. While some canonical variants may indeed contribute to resistance in specific clades, their effects may be masked in population-based analyses due to strong correlations with genetic ancestry. These findings challenge the view that azole resistance is primarily driven by changes in drug efflux or target availability, highlighting instead the diverse contributions of mutations within a polygenic framework.

Heritability estimates indicated that these phenotypes were all highly heritable. Furthermore, the fact that *h^2^_SNP_* was much greater than the variance explained by genome-wide significant associations suggests a highly polygenic architecture to these traits, and that, as sample sizes grow, additional smaller effect loci will be identified. Thus, other aspects to the genetic etiology of azole response phenotypes are yet to be characterized. Notably, genetic correlations revealed a significant but partial overlap between susceptibility and tolerance, while heteroresistance displayed antagonistic relationships with susceptibility, highlighting potential trade-offs in evolutionary strategies for azole adaptation. This antagonistic relationship is further supported by opposing GWAS signals in a shared region on Chr5. Thus, heteroresistance appears to occupy a distinct position in the evolutionary landscape of azole adaptation, phenotypically overlapping with both susceptibility and tolerance while maintaining a unique genetic architecture.

At the population level, we observe that chromosome copy number variation also influences azole responses, particularly susceptibility, supporting the idea that dosage-sensitive mechanisms impact drug adaptation. Indeed, aneuploidy has been previously associated with strains adapting to azole exposure. For example, an isochromosome 5 underlies resistance (62), and ChrR trisomy was detected in tolerant isolates (63). Importantly, we find that karyotypic variation itself, rather than specific chromosome gains, enhances fitness under drug pressure. Although chromosomal aneuploidy can contribute to phenotypic variation and could, in principle, be modeled as a variable in GWAS, we did not include it in our current analysis. Aneuploidies often exert broad, pleiotropic effects on cellular physiology and genome stability that do not behave like traditional Mendelian loci, potentially violating key assumptions of GWAS models and complicating the interpretation of association signals.

Our functional validation of GWAS-prioritized genes revealed that natural variation in diverse cellular processes, including cell wall integrity, endocytosis, DNA repair, and telomere maintenance, can strongly modulate azole responses. Because these mutant analyses were conducted in a single genetic background (SC5314), the observed effects may underestimate the full phenotypic potential in clinical populations. Nonetheless, these experiments provide key links between GWAS associations and biological mechanisms, uncovering novel regulators of azole responses. Future work introducing the precise GWAS-identified SNPs into diverse genetic backgrounds will better model natural variation. Additionally, the absence of strong phenotypic effects from any single gene in this analysis suggests that the observed phenotype may arise from the combined influence of multiple genes, consistent with the polygenic signals detected in our genetic analyses.

Together, these results reveal the complex genetic landscape of azole adaptation in *C. albicans*, where polygenic mechanisms, aneuploidy, and evolutionary trade-offs drive resistance, tolerance, and heteroresistance. Notably, the identification of key associations with DNA maintenance genes uncovers a previously underappreciated link between genome stability pathways and antifungal responses. The convergence of these adaptive mechanisms highlights the need for combination therapies that simultaneously target drug efflux, stress pathways, and genome integrity to outpace fungal evolutionary capacity and prevent treatment failure.

## MATERIALS AND METHODS

In this study, we assessed 557 total isolates for azole susceptibility, tolerance, and heteroresistance, then utilized whole genome sequence (WGS) data to perform a GWAS to identify loci associated with each phenotype. We used genome-wide data to estimate the genetic correlations of these phenotypes and used phylogenetic approaches to investigate the evolutionary history of resistance, heteroresistance, and tolerance in these lineages.

### Isolate sampling

The 557 *C. albicans* isolates were gathered from diverse environments, including from food spoilage and a starling (*Sturnidae*), as well as sites of human infection and colonization (Fig. S1A and Table S1). The isolates were compiled from four separate collections to maximize genetic and phenotypic diversity. We included 192 isolates spanning all major *C. albicans* lineages; these comprised 145 strains from published work (30, 64) and 47 new isolates (Table S1). We also incorporated 7 isolates from individuals with nonpersistent invasive infections and 56 longitudinal isolates from 12 individuals with persistent invasive infections, previously used to investigate azole tolerance (7). We included 40 longitudinal isolates from HIV patients undergoing fluconazole treatment for oropharyngeal candidiasis that developed azole resistance during treatment (3), 269 single and longitudinal isolates from patients with oropharyngeal candidiasis (gift of Prof. Dominique Sanglard, University of Lausanne), and 2 isolates from Hirakawa et al (65). Finally, we included 8 *Candida africana* isolates (30) which were used as a phylogenetic outgroup.

### C. albicans growth

*C. albicans* strains were grown overnight in liquid YPD medium (2% bacto-peptone, 1% yeast extract, and 2% dextrose (filter-sterilized)) at 30°C with continuous shaking at 200 rpm. The cell density of each culture was determined by measuring the optical density of diluted cultures at 600 nm (OD_600_) in sterile water using a Biotek Epoch 2 microplate reader (Agilent Technologies). The cultures were subsequently diluted to the desired concentrations in sterile water.

### Phenotyping fluconazole susceptibility and tolerance

We quantified fluconazole susceptibility for each strain by measuring the minimal inhibitory concentration (MIC_50_), following previous studies (66). MIC_50_ testing was performed using broth microdilution assays in 96-well plates. Fluconazole (PHR1160-1G, Sigma Aldrich) was serially diluted (1 in 2) in liquid YPD to 12 final concentrations ranging from 0 to 256 μg/mL. Each well contained 125 μL liquid YPD volume with *C. albicans* cells at a concentration of 2 x 10^5^ cells/mL. Plates were incubated at 30°C with shaking (200 rpm) and cell densities (OD_600_) were measured at 0, 24, and 48 h using a Biotek Epoch2 spectrophotometer. The plates were examined for contamination after each incubation period. MIC_50_ values were determined after 24 h of growth by identifying the drug concentration leading to ≤ 50% growth relative to growth without fluconazole (FLC). Following Alabi et al (2023), we then measured tolerance as supra-MIC growth (SMG), at 48 h. SMG corresponds to the average growth in wells above MIC_50_ values compared to those without the drug. Thus, an SMG value of 1 indicates that the strain presents equal cell density in wells with drug concentrations above the MIC_50_ value as in those without the drug, while intermediate values (e.g., SMG of 0.5) indicate that the strain’s growth remains impaired by the drug at 48 h after inoculation. Tolerance could not be measured for isolates with MIC_50_ greater than 128 μg/ml, resulting in 485 isolates with measurable tolerance. All assays were performed with at least three biological replicates; averaged values are included in the Table S2.

### Phenotyping fluconazole heteroresistance

Following Gautier et al. (11), we conducted population analysis profiling (PAP) assays to quantify heteroresistance across the 557 strains. Cells were grown overnight in YPD medium, and cell densities were adjusted to 10⁶ cells/mL, from which three subsequent dilutions were prepared: 10⁵, 10⁴, and 10³ cells/mL. Three droplets of 5 µL from these cell suspensions were plated onto YPD agar supplemented with either 0 or 128 μg/mL FLC. The plates were incubated at 30°C and imaged at 48 h to determine the number of non-confluent colonies from the different cell dilutions. Heteroresistance was calculated as the number of non-confluent colonies at 128 μg/mL FLC divided by the number of such colonies in the absence of FLC. Highly heteroresistant strains will thus be present as a high fraction of the overall population. Since heteroresistance is calculated using FLC concentrations that are 10-fold higher than the MIC _50_, it could not be determined for isolates with an MIC_50_ greater than 32 μg/ml, resulting in 422 isolates with measurable heteroresistance. All assays were performed with at least three biological replicates; averaged values are included in Table S2.

### Mutant analyses

The susceptibility, tolerance, and heteroresistance of genetic mutants were measured as described above. For the overexpression strains, the YPD plates were supplemented with anhydrotetracycline hydrochloride (ATC) (10792081, Thermo Scientific Acros) at 3 µg/mL and supplemented with 1 mM of FeCl₃ (220299-250g, Sigma Aldrich) to mitigate the ATC effects on iron chelation. For the overexpression strains, the MIC_50_/SMG/heteroresistance levels of the mutant strains were calculated by dividing each phenotype value in the presence of ATC relative to the corresponding value without ATC and comparing to the same ratio to the wild type (parent) isolate.

### Genomic extraction, Illumina sequencing, and variant calling

For the *C. albicans* isolates without published genomes, the isolates were streaked from –80° C stocks on YPD plates and incubated at 30° C overnight. Single colonies were selected and resuspended in 5 ml YPD medium for overnight incubation at 30°C with continuous shaking at 200 rpm. The cultures were centrifuged and washed with sterile PBS, and the DNA was extracted from the resulting cell pellets using the Qiagen Genomic DNA kit utilizing the QIAGEN Genomic-tip20/G. DNA extractions were performed according to the manufacturer’s instructions. Sequencing libraries were prepared by SeqCenter (Pittsburgh, PA) using the Illumina DNA Prep and sequenced on an Illumina NovaSeq 6000. Sequencing depth is listed in the Table S1.

The genome sequences and General Feature Files for the SC5314 reference genome (haplotype A, version A22-s07-m01-r130) were downloaded from the *Candida* Genome Database (http://www.candidagenome.org/). Reads were aligned to the SC5314 reference genome (haplotype A chromosomes) using Minimap2 version 2.17 (67). Sequence Alignment/Map tools (SAMtools) v1.10 (r783) and Picard tools version 2.23.3 (http://broadinstitute.github.io/picard) were used to filter, sort, and convert the SAM files. SNPs were called using the Genome Analysis Toolkit (GATK) version 3.6 (68, 69), according to the GATK Best Practices. SNPs were filtered using the following parameters: VariantFiltration, QualByDepth (QD) < 2.0, LowQD, ReadPosRankSum < −8.0, LowRankSum, FisherStrand (FS) > 60.0, HightFS, MQRankSum < 12.5, MQRankSum, MQ < 40.0, LowMQ, HaplotypeScore > 13.0. Coverage levels were calculated using GATK. A dataset of 647249 confident SNPs across the isolates was created. Besides passing GATK’s filters, we also classified heterozygous positions as those with an allelic ratio of the number of alternative allele reads/total number of reads comprised between 20% and 80% and homozygous positions as those with an allelic ratio of the number of alternative allele reads/total number of reads of >80% and <20%. Plink files were generated with PLINK 2.0 (www.cog-genomics.org/plink/2.0/) (70).

### Maximum likelihood phylogenetic inference

We estimated the phylogenetic relationships among the samples through maximum likelihood (ML) analysis of the SNP alignment with 565 taxa (including 8 *Candida africana* strains as outgroups) and 1,294,498 sites. The ML tree for the concatenated dataset was estimated with RAxML-NG v.1.0.1 using the General Time Reversible (GTR) + G + I model of nucleotide substitution with 100 bootstrap replicates to assess support (71). Isolation source, clade identification, and azole phenotypes were mapped and visualized using iTOL (72).

### Quality control and SNP filtering

Using Plink v1.9 (73), we conducted quality control on 557 *C. albicans* isolates and 628,048 SNPs, evaluating minor allele frequency (MAF), missing genotype rates, heterozygosity, LD, and Hardy-Weinberg Equilibrium (HWE). A total of 228,346 SNPs violated the HWE threshold (p < 1e-6), and 369,698 SNPs had a MAF < 0.001. For population structure analyses, we filtered the dataset down to 145129 high-quality, independent SNPs (MAF > 0.05, HWE *P* > 1e-6). To evaluate the impact of rare variant inclusion on test statistic inflation, we performed GWAS across multiple MAF) bins. We observed that incorporating increasingly rare variants resulted in a progressive decrease in the genomic inflation factor (λ), indicating reduced statistical power and potential inflation control issues at lower MAF thresholds.

### Genome-wide association testing

The GWAS analyses for each phenotype (FLC susceptibility, tolerance, and heteroresistance) were conducted separately using MIC_50_ scores, SMG values, and PAP fractions, respectively. These analyses were performed using GCTA v1.94.1 (51) with the MLMA-LOCO method (-- mlma-loco). MLMA-LOCO is a mixed linear model analysis that excludes the chromosome, on which the candidate SNP is located, from calculating the genetic relationship matrix (GRM) in GCTA v. 1.94.1. The GRM used in these models was derived from stringently filtered genotype array data using Plink v1.9 (**-**-maf 0.05 --hwe 1e-6 --geno 0.1 --indep-pairwise 50 5 0.2; n = 145,129/628,048 remaining variants) (73). All individuals included in the analyses have a pair-wise genetic similarity < 0.025. In GCTA, we performed an eigen-decomposition of the GRM (--pca 2) to calculate principal component axes (PCs) to include as covariates in the MLMA to correct for population structure. We calculated λ as the ratio of the median observed chi-square test statistic to the expected median of a chi-square distribution with 1 degree of freedom. A λ score close to one indicates that there is no inflation of test statistics and that population structure is accounted for in the model. Across all three phenotypes, we retained only 2 PCs to be included as covariates in the MLMA model, as including more than two overcorrected for population substructure (λ >1). Manhattan and QQ plots were visualized using summary statistics from the MLMA-LOCO analysis in the R package “CMPLOT” (72).

P-value thresholds (α) for each phenotype were determined through permutation testing. Phenotype data were permuted 1,000 times, and an MLMA-LOCO analysis was performed on each permuted dataset. For each permuted phenotype dataset, we repeated the MLMA-LOCO, then extracted the minimum *P* value. The 5th percentile of this minimum *P* value distribution, derived from 1,000 null phenotype GWASs, was used as a threshold for genome-wide significance. The significant *stringent* thresholds applied were as follows: α_tol_ = 1.12E^-05^, α_sus_= 3.57E^-06^, and α_het_ = 1.88E^-05^. We also set a *relaxed* alpha threshold one order of magnitude higher (less strict) than the calculated alpha for each phenotype to examine additional weaker associations that included candidate loci. To account for the differing scales of phenotypic measurements, we calculated the proportion of variance explained (PVE) by each SNP using the formula: PVE = 2pqβ/VP where *p* is the frequency of one allele, *q* =1−*p* is the frequency of the alternate allele, and β represents the estimated effect size of the SNP. Here, *V_P_* is the total phenotypic variance for each phenotype. To reduce redundancy due to LD, we performed COJO (conditional and joint analysis) in regions with clusters of associations, but not genome-wide. Therefore, some LD inflation may remain in the total PVE estimate. Additionally, as these estimates were based on the discovery sample, they are likely subject to the winner’s curse, and the reported total PVE may be an overestimate of the true variance explained in an independent sample.

### COJO analysis

We used the Conditional and Joint (COJO) Analysis in GCTA v. 1.94.1 (--cojo-slct --cojo-p 1e-4) to identify independent genetic signals from our GWAS summary statistics in regions with a high number of significantly associated SNPs (44). We generated a LD matrix for pairwise SNPs within these regions using PLINK v1.9. This methodology enabled us to differentiate between multiple association signals within shared regions, specifically on Chr5 and Chr3. We also created LD matrices for pairwise SNPs in these regions using PLINK v1.9.

### Heritability and genetic correlation by phenotype

In GCTA v. 1.94.1 (51), a Genomic-Relatedness-based Restricted Maximum Likelihood (GREML) analysis was conducted to estimate variance components and SNP-heritability (*h^2^_SNP_*), the proportion of phenotypic variance explained by markers in the GRM. Estimates of *h^2^_SNP_* were calculated (GCTA command: --grm pca_output –reml) for each phenotype separately (susceptibility, tolerance, and heteroresistance) using 2 PC vectors and the GRM to account for population structure. A bivariate GREML Analysis was conducted in GCTA v. 1.94.1 (51) to determine the shared genetic architecture and the genetic covariance structure between traits. We estimated SNP-heritability for each chromosome using chromosome-specific GRMs in GCTA and assessed the relationship between chromosome length and SNP-heritability using a Spearman correlation test. Pairwise genetic correlation (*rG*) (54) between each phenotype was estimated with residual covariance using the --reml-bivar flag.

### Mapping the phylogenetic distribution of significant loci

For each significantly associated locus, we extracted the carrier status from PLINK version 1.9 using the --extract flag followed by the relevant SNP information in text format and then recoded to designate which isolates were homozygous or heterozygous. Carrier status was then converted into a heatmap matrix and aligned to the previously estimated ML phylogeny using the R package “ggtree” (Yu et al. 2017).

### Data availability

The sequence data from this study have been submitted to NCBI Sequence Read Archive under BioProject ID PRJNA1264048 (http://www.ncbi.nlm.nih.gov/bioproject). Code for this project can be found at https://github.com/users/kyleschutz/projects/3.

## ACKNOWLEDGEMENTS

We thank Prof Dominique Sanglard (University of Lausanne) for the gift of *C. albicans* strains. Work in the laboratory of CdE was supported by the Agence Nationale de Recherche (ANR-10-LABX-62-IBEID). LE was supported by funding from Hevolution/AFAR HEV∼NI23013 and NIA RO1AG046392. SM was supported by funding from NSF DEB-1553114. IVE was supported by funding from the Agence Nationale de Recherche (JCJC GENOMEHET) and the Institut Pasteur.

