## Supplemental Figures for "Polygenetic Determinants of Azole Resistance, Tolerance, and Heteroresistance in *Candida albicans*"

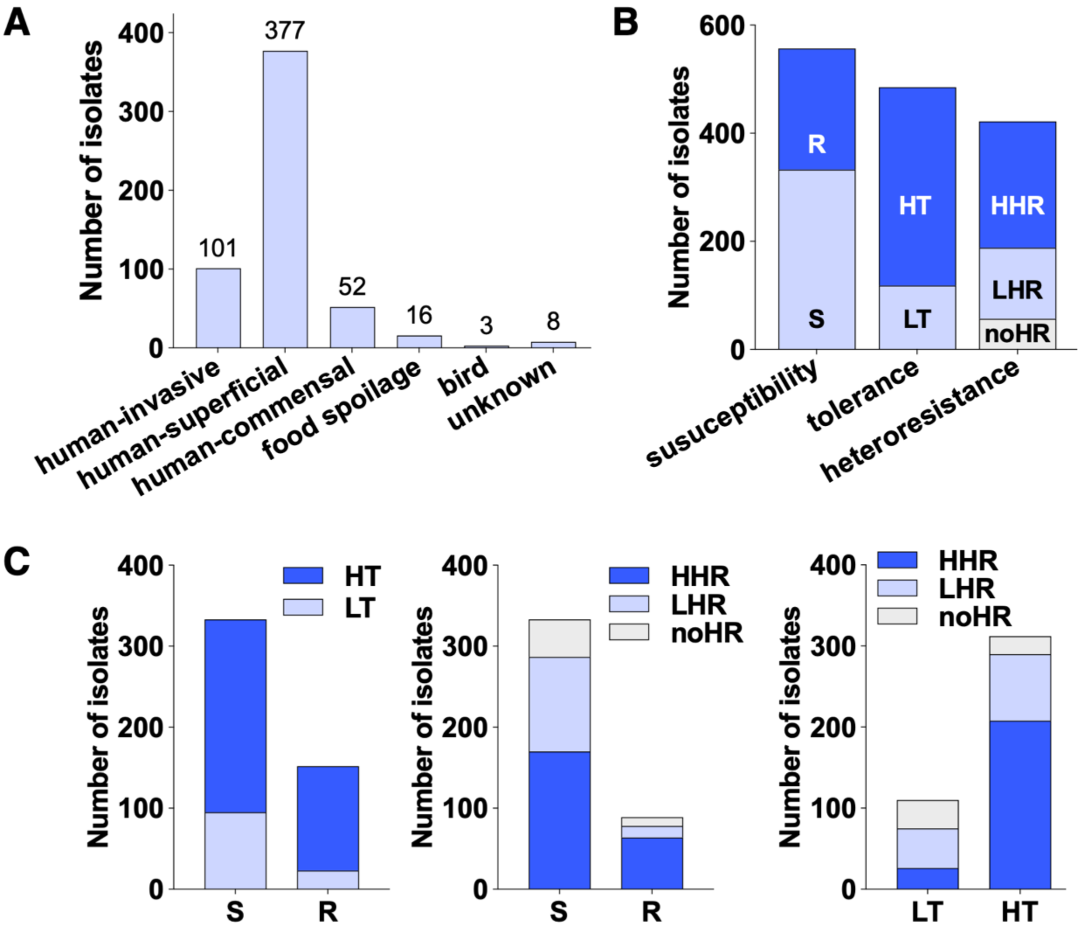


**Figure S1. Strain characteristics and phenotype distributions.** (A) Distribution of *C. albicans* strains across isolation sources. (B) Number of *C. albicans* isolates showing susceptibility (S), resistance (R), low tolerance (LT), high tolerance (HT), no heteroresistance (noHR), low heteroresistance (LHR), or high heteroresistance (HHR). Tolerance and heteroresistance could only be measured in isolates with $MIC_{50}$ <128 µg/ml and $MIC_{50}$ <64 µg/ml, respectively. (C) Panels show the number of susceptible and resistant strains showing high vs. low tolerance and high vs. low vs. no heteroresistance, and the number of high and low tolerance isolates showing the three heteroresistance categories.

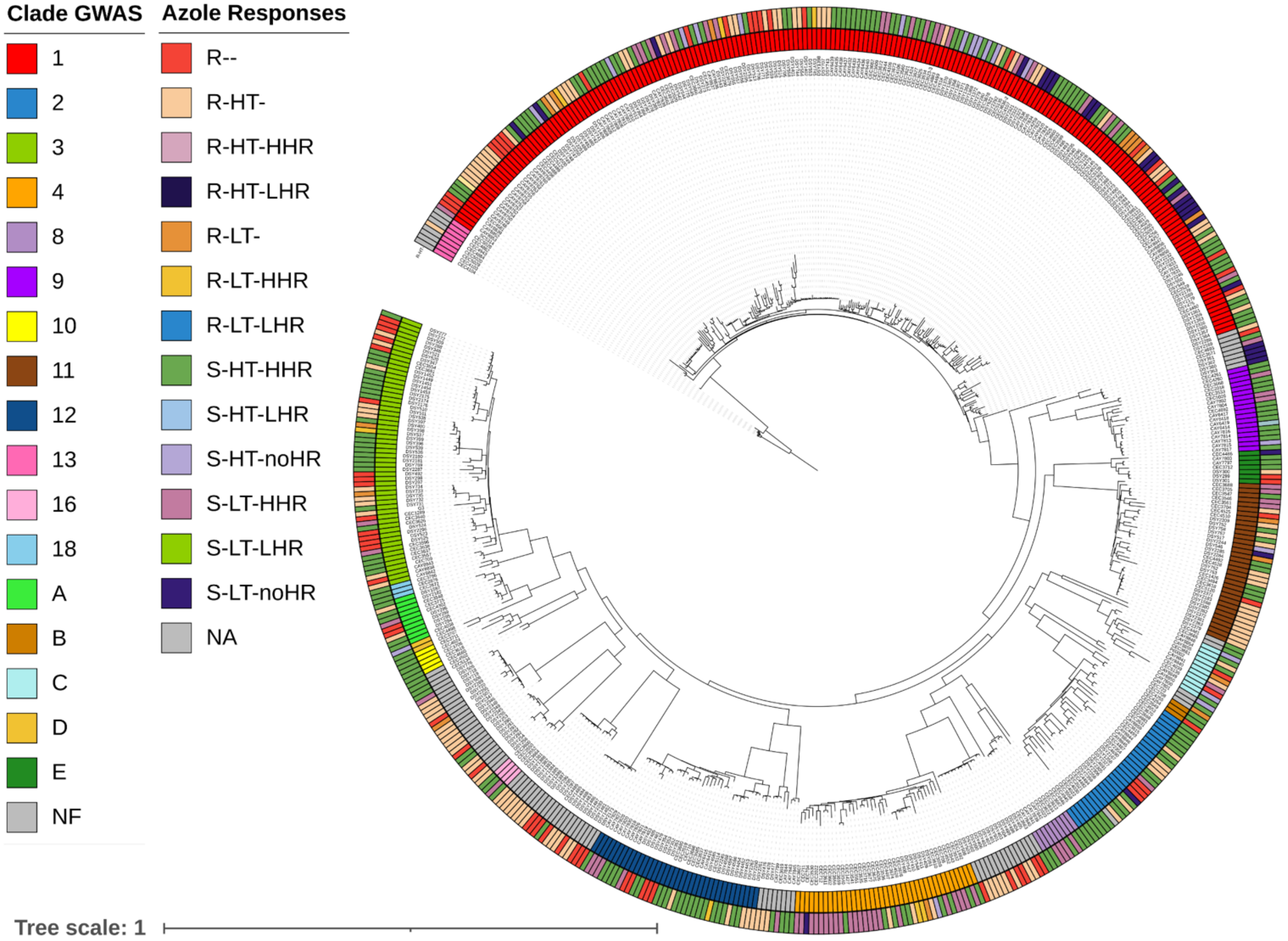


**Figure S2. Phylogenetic distribution of phenotypes.** Distribution of drug response phenotypes across the estimated maximum likelihood (ML) phylogeny with binned azole response phenotypes values mapped to tree branches in the outer ring (susceptibility (S), resistance (R), low tolerance (LT), high tolerance (HT), no heteroresistance (noHR), low heteroresistance (LHR), and high heteroresistance (HHR)) and phylogenetic clades on the inner ring.

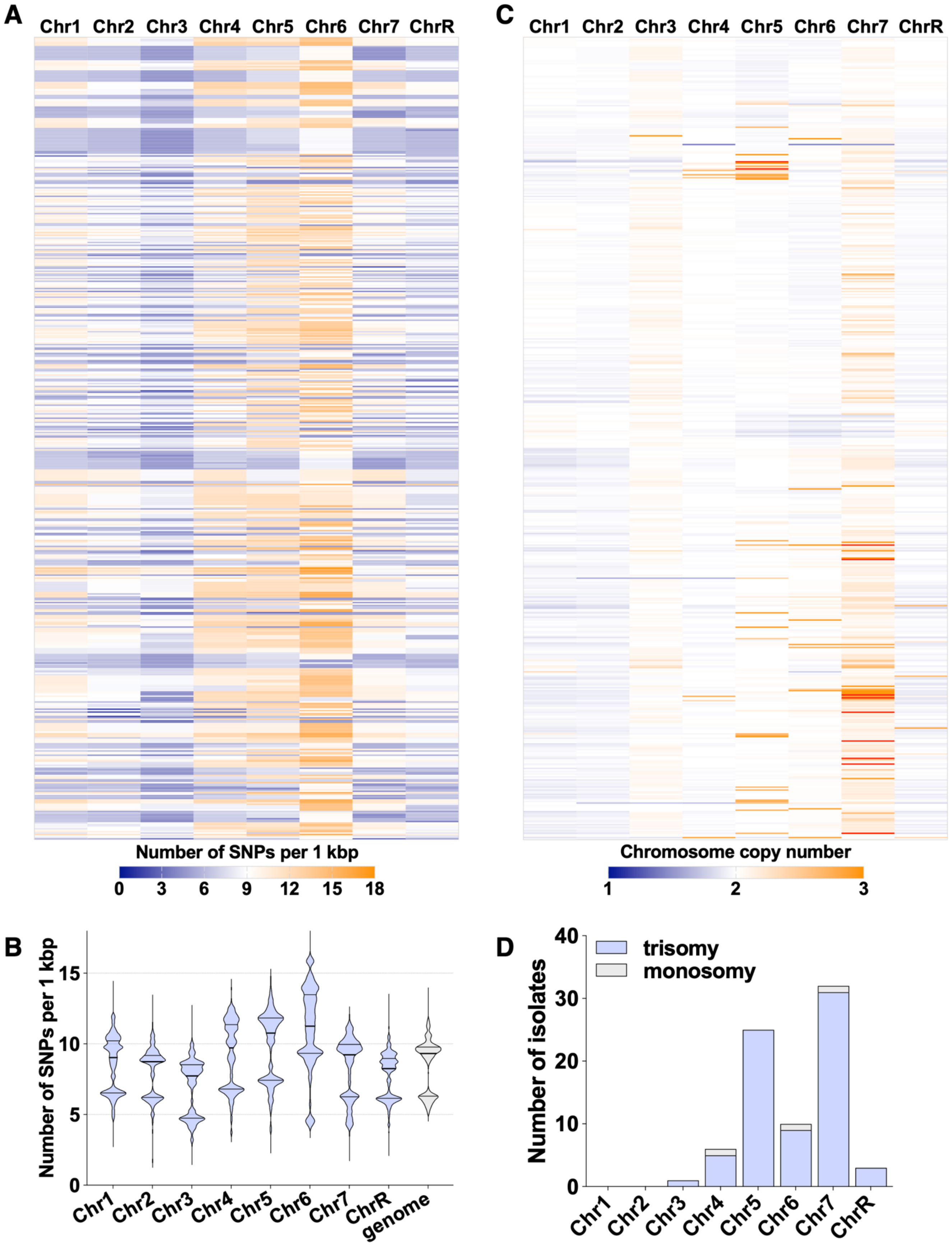
A
Chr1 Chr2 Chr3 Chr4 Chr5 Chr6 Chr7 ChrR
Number of SNPs per 1 kbp
0 3 6 9 12 15 18
C
Chr1 Chr2 Chr3 Chr4 Chr5 Chr6 Chr7 ChrR
Chromosome copy number
1 2 3
B
Number of SNPs per 1 kbp
0 5 10 15
Chr1 Chr2 Chr3 Chr4 Chr5 Chr6 Chr7 ChrR genome
D
Number of isolates
0 10 20 30 40
trisomy
monosomy
Chr1 Chr2 Chr3 Chr4 Chr5 Chr6 Chr7 ChrR

**Figure S3. Landscape of genetic variation across the 557 *C. albicans* isolates.** (A) SNP density per chromosome. (B) Average number of SNPs per 1 kbp for each chromosome. (C) Chromosomal copy number variation across isolates. (D) Number of isolates with chromosomal copy number variants (trisomy or monosomy) on each chromosome.

**A**

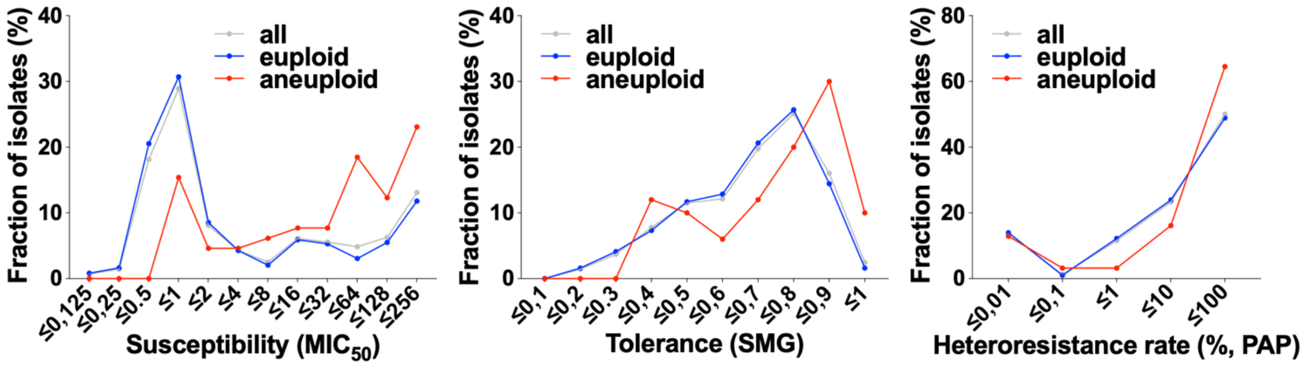


**B**

| Statistic | Karyotype | Susceptibility (MIC$_{50}$) | Tolerance (SMG) | Heteroresistance (%) |
|---|---|---|---|---|
| average | aneuploid | 88.76 | 0.69 | 39.39 |
| | euploid | 41.54 | 0.63 | 32.59 |
| median | aneuploid | 53.33 | 0.75 | 40.97 |
| | euploid | 41.54 | 0.63 | 32.59 |

**C**

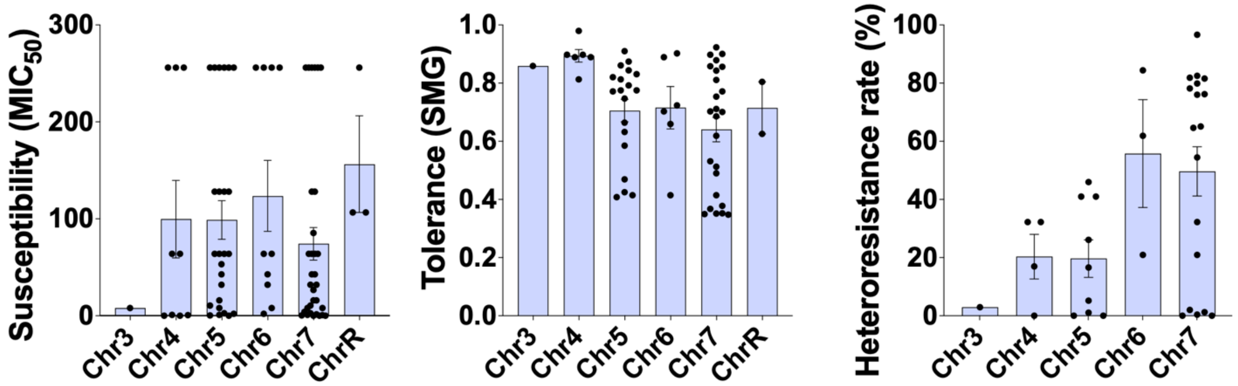


**Figure S4. Karyotype-phenotype associations.** (A) Distribution of susceptibility (MIC$_{50}$), tolerance (SMG), and heteroresistance rates (%) across euploid (blue), aneuploid (red), or all (grey) isolates. (B) Phenotype distributions in euploid vs. aneuploid isolates, showing average and median values across the three drug responses. (C) Phenotype levels (susceptibility, tolerance, and heteroresistance rate) for aneuploids of each chromosome reveal a lack of association between specific aneuploids and drug response levels.

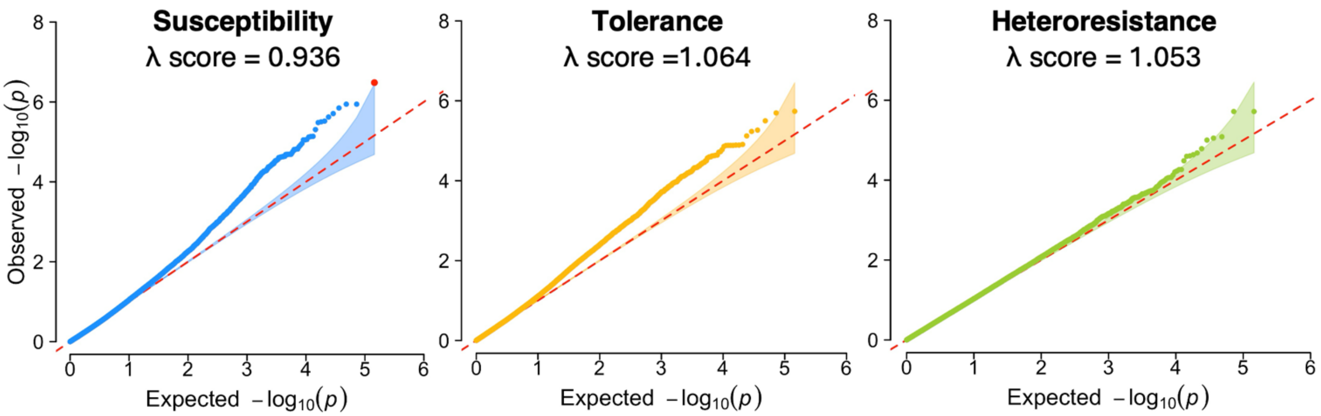


**Figure S5. Quantile-quantile (QQ) plots for GWAS analysis.** QQ plots indicate the fit of the data to the MLMA (observed vs. expected −log10(*P*)) for (A) susceptibility, (B) tolerance, and (C) heteroresistance. λ (genomic inflation) scores are also included for each drug response.

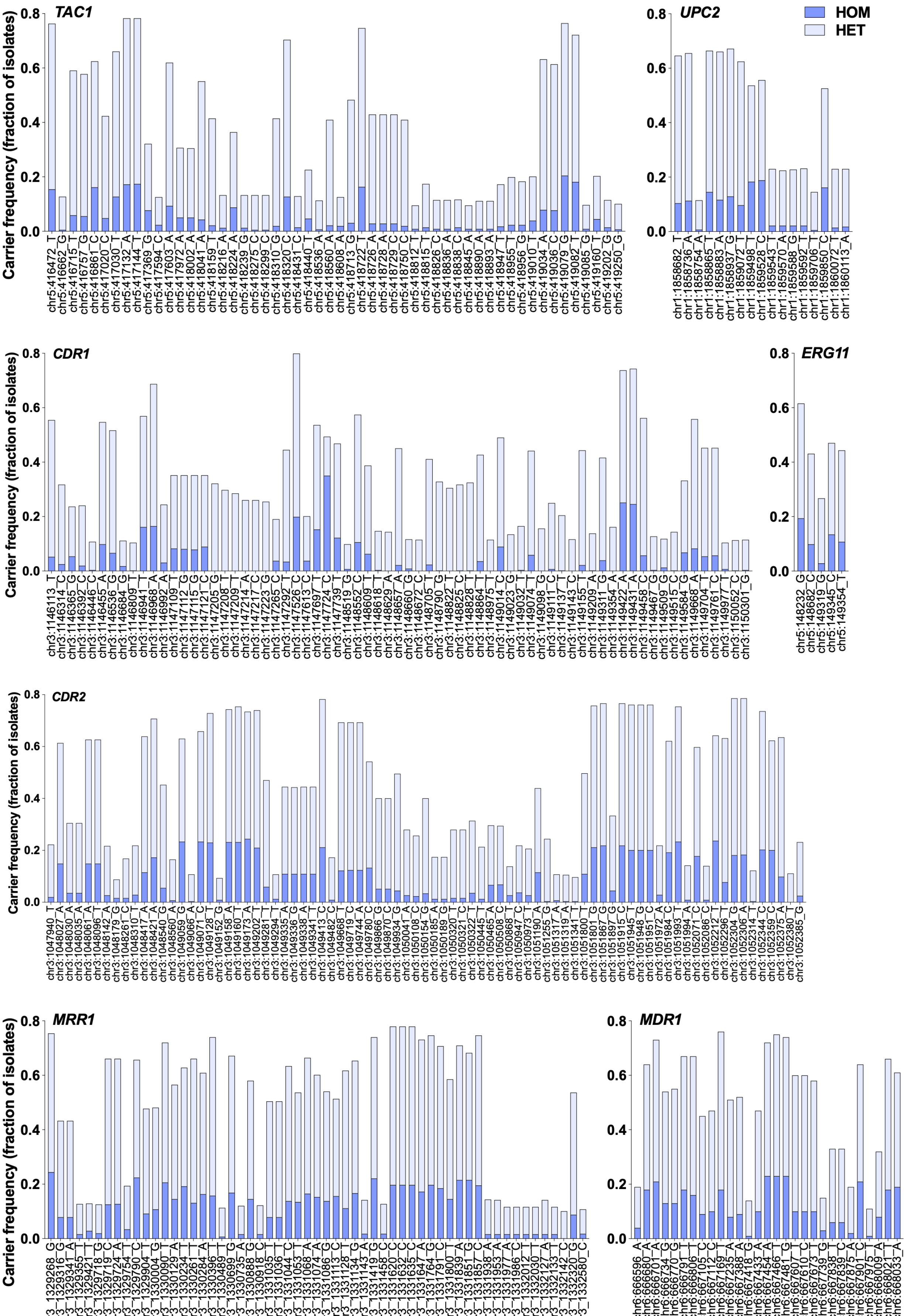

TAC1
UPC2
HOM
HET
CDR1
ERG11
CDR2
MRR1
MDR1
Carrier frequency (fraction of isolates)

**Figure S6. Allelic variation in genes previously associated with *C. albicans* azole resistance.** Carrier frequencies of heterozygous and homozygous SNPs of canonical resistance genes (*CDR1, CDR2, ERG11, MDR1, MRR1, TAC1,* and *UPC2*) across the 557 *C. albicans* isolates.

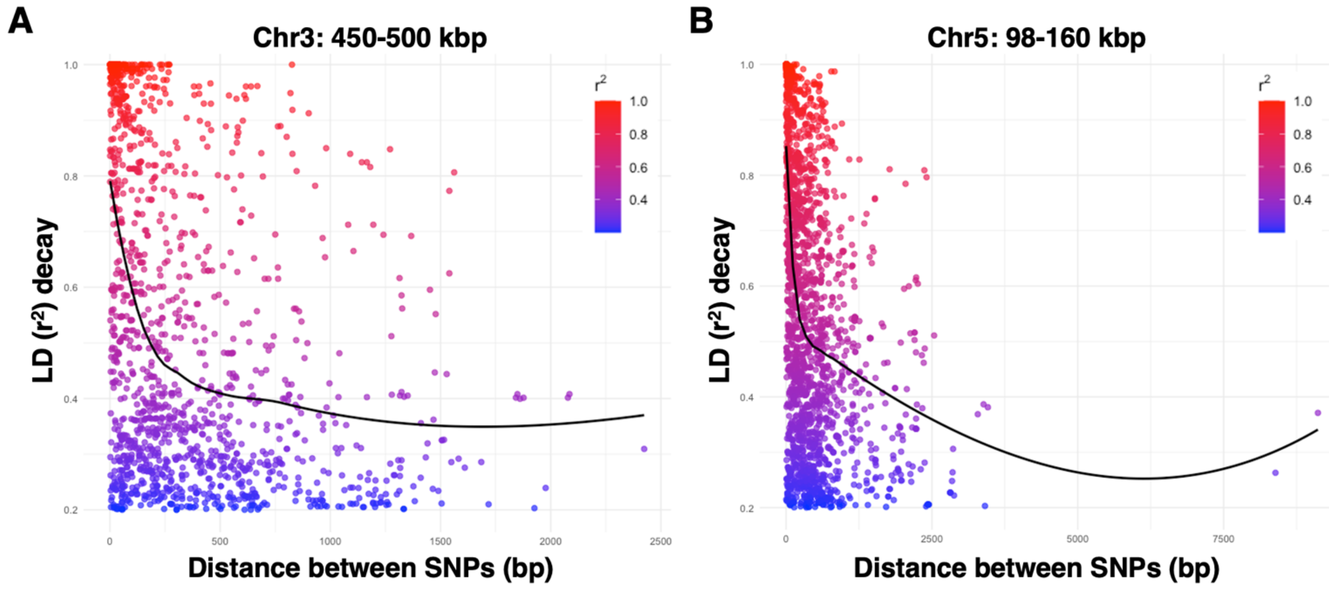


**Figure S7. Linkage disequilibrium (LD) in Chr3 and Chr5 regions.** LD decay plots showing the distribution of pairwise $r^2$ values as a function of genomic distance for SNPs located in regions of Chr3 (A) and Chr5 (B) included in the COJO analysis.

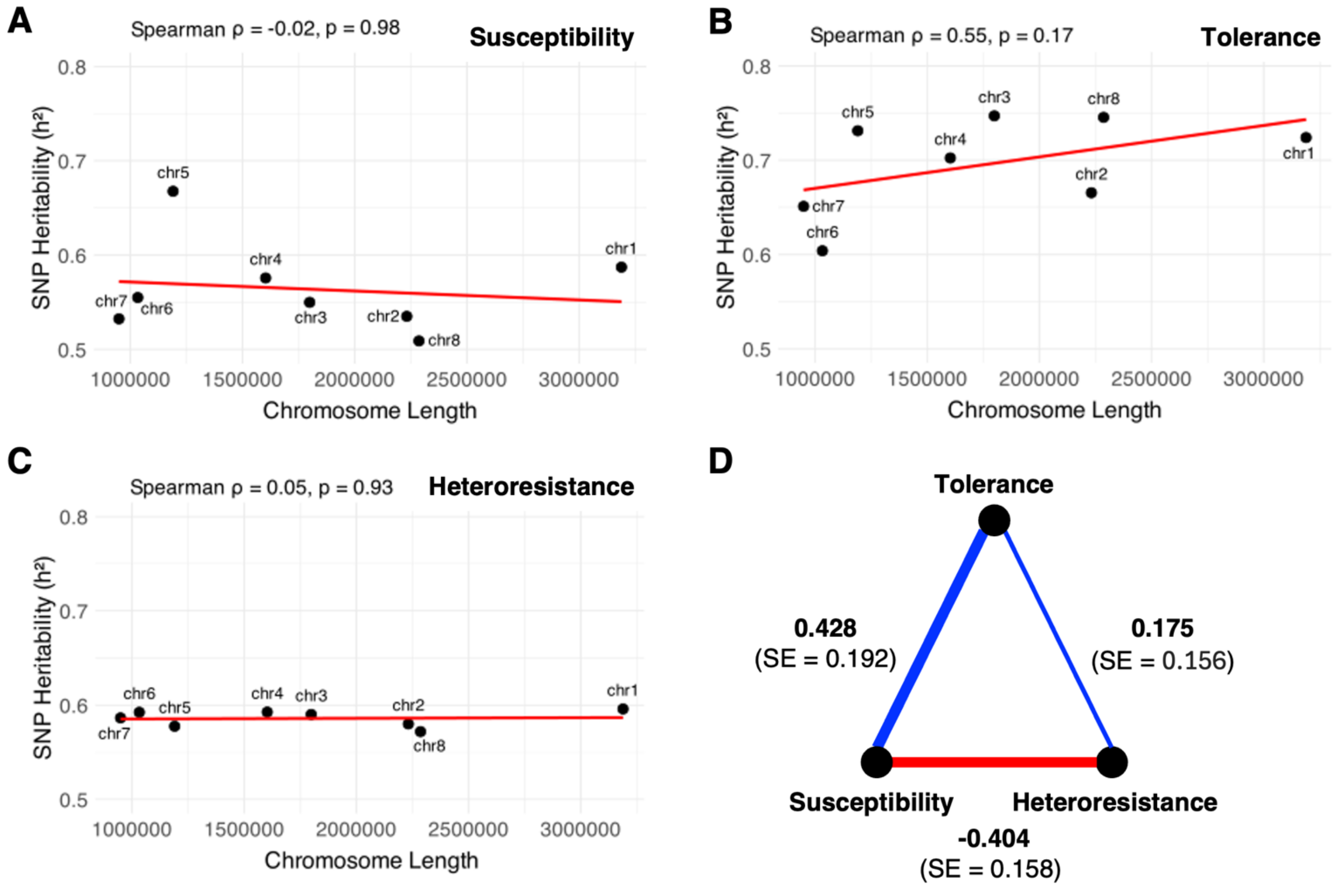


**Figure S8.** SNP-heritability ($h^2_{SNP}$) for susceptibility (A), tolerance (B), and heteroresistance (C), calculated for each *C. albicans* chromosome. Correlations between chromosome size and heritability were determined using a Spearman correlation test. (D) Genetic correlations and standard error (rG(SE)) between susceptibility, tolerance, and heteroresistance using a bivariate GREML analysis. Positive values of rG indicate that genetic influences on one trait also influence the other trait in the same direction, while negative values suggest opposing effects. These correlations are derived from the genome-wide relationship matrix (GRM) based on SNP data.

**Figure Legends for Supplementary Tables**

**Table S1.** *Candida* isolates and mutant strains used in this study. The table includes the source of isolation, sequencing details, and NCBI repository information for each GWAS isolate.

**Table S2.** Susceptibility, tolerance, and heteroresistance levels across the 557 *C. albicans* isolates. Phenotype bins were defined as S (susceptible), R (resistant), HT (high tolerance), LT (low tolerance), HHR (high heteroresistance), LHR (low heteroresistance), noHR (no heteroresistance).

**Table S3.** Stringent and relaxed GWAS loci identified for azole susceptibility, tolerance, and heteroresistance. Statistics regarding the proportion of explained variance (PVE) for different types of SNPs and top genes are also included.

**Table S4.** True associations recovered through COJO analysis.

**Table S5.** Phenotypic variance and covariance using GREML analyses.

**Table S6.** SNPs with antagonistic effects on different azole drug responses.

**Table S7.** Susceptibility ($MIC_{50}$) and tolerance (SMG) levels for mutant strains tested in heteroresistance assays, shown relative to the corresponding parental strains.
